# Genomic signatures of adaptation in native lizards exposed to human-introduced fire ants

**DOI:** 10.1101/2023.09.24.559217

**Authors:** Braulio A. Assis, Alexis P. Sullivan, Stephanie Marciniak, Christina M. Bergey, Vanessa Garcia, Zachary A. Szpiech, Tracy Langkilde, George H. Perry

## Abstract

Understanding the process of genetic adaptation in response to human-mediated ecological change will help elucidate the eco-evolutionary impacts of human activity. Red fire ants (*Solenopsis invicta*) spread across Southeastern USA since their accidental introduction via Port Mobile, Alabama in the 1930s, serving today as both novel venomous predator and novel toxic prey to native eastern fence lizards (*Sceloporus undulatus*). To identify potential signatures of genetic adaptation in lizards to invasive fire ants, we generated whole genome sequencing data from 420 native fence lizards sampled across three populations, two of which had not been invaded by fire ants (in Tennessee and Arkansas) and one which had been invaded for ∼70 years (Alabama). We detected signatures of positive selection exclusive to the exposed Alabama population for genetic variants overlapping genes related to the membrane attack complex of the complement immune system, growth factor pathways, and morphological development. Prior work identified a relationship between increased lizard survival of fire ant attack and longer hind limbs, which lizards use to remove ants from their bodies. Furthermore, we conducted a genome-wide association study with 381 Alabama lizards to identify 24 hind limb length-associated genetic loci. For two loci, positive-effect alleles occur in high frequency and overlap genomic regions that are highly differentiated from the populations naïve to fire ants. Collectively, these findings represent plausible genetic adaptations in response to fire ant invasion, whereby morphological differentiation may increase survival against swarming ants and altered immune responses may allow the exploitation of a novel, toxic food resource.

**Significance statement:** Human activity can force interactions between species from distinct ecological backgrounds. These interactions can consequently impose novel selective pressures on endemic populations via predation or disruption of ecological niches through community-wide effects. While some endemic taxa have been able to adapt biologically to these disruptions, we do not have a full understanding of the underlying genetic processes that may allow it. Here we identify genomic signatures of recent adaptation nearby genes involved in morphological and immunological processes in native fence lizards that are consistent with pressures imposed by the venomous, predatory fire ants introduced by humans. These signatures are largely absent from lizard populations that are naïve to fire ants.

## Introduction

Endemic species worldwide face rapid environmental change resulting from various types of human activity. For example, human-mediated translocation of species into new environments promotes novel ecological interactions, often with detrimental effects (Saul and Jeschke 2015). The consequences can be dire: species introductions have been identified as an underlying cause of at least 170 animal extinctions (Clavero and García-Berthou 2005). In some invasive-endemic interaction cases, standing genetic variation may provide endemic populations with the raw materials needed for quick adaptative responses (Barrett and Schluter 2008). Yet even then, such adaptations can induce cascading effects on the broader ecosystem (Mooney and Cleland 2001; Schlaepfer et al. 2005; Strauss et al. 2006; Hale et al. 2016). Therefore, understanding the process of genetic adaptation to novel species interactions will better inform us of the scope of potential ecological impacts related to human behavior.

In the 1930s, the red imported fire ant, *Solenopsis invicta*, was accidentally transported, by humans, from South America (presumably northeastern Argentina) to Port Mobile, Alabama, in the United States of America (Ascunce et al. 2011). Since then, fire ants have established and steadily expanded their range in the Southeastern USA, with both economic and public health impacts arising from their potent venom and aggressive nature (Gruber et al. 2022). Furthermore, ecological impacts stemming from fire ant invasion are marked and diverse. In the United States, fire ants outcompete and displace native ants, with cascading effects on the broader invertebrate community (Porter and Savignano 1990; Morrison 2002; Roeder et al. 2021). Experimental studies have confirmed that fire ants directly and indirectly impact various endemic small vertebrates. Specifically, predation by fire ants led to a reduction of up to 66% of *Ambystoma* salamander populations within 48h (Todd et al. 2008), whereas fire ant suppression led to significant increases in small vertebrate abundance (Stahlschmidt et al. 2018). Meanwhile, fire ant disruption of arthropod communities led to a 10% decrease in the number of eastern bluebird fledglings and to the displacement of adult birds (Ligon et al. 2011).

Among impacted vertebrates, the eastern fence lizard *Sceloporus undulatus* has been an important model system for studying the ecological impacts of this fire ant introduction. First, despite their toxicity, fire ants have become a novel prey item in the fence lizard’s diet (Robbins et al. 2013). Second, fire ants are a novel predator of fence lizards: swarming worker ants can envenomate and kill juvenile and adult individuals (Langkilde 2009), as well as prey on their eggs (Thawley and Langkilde 2016). The predatory impact is likely very high: experimental removal of fire ants from enclosures increased fence lizard recruitment by ∼60% (Darracq et al. 2017), while lizard hatchling survival is negatively associated with fire ant mound density (Gifford et al. 2017). Even lizards that initially survive fire ant encounters experience a 20% increase in mortality rate over the 11 weeks post-exposure relative to unexposed lizards (Langkilde and Freidenfelds 2010). Yet, fence lizards remain abundant in fire ant-invaded habitats, prompting behavioral, ecological, and evolutionary investigations into the underlying mechanisms.

Exposure to fire ant venom induce wide-ranged immunological responses in fence lizards (Tylan et al. 2020; Tylan et al. 2023). Prior to venom exposure, however, behavioral defenses are also employed by fence lizards against swarming fire ants. These defenses include body-twitching – using their hind legs to directly remove ants with a flicking motion – and fleeing from the attack (Langkilde 2009). In behavioral trials, the success of these tactics was positively linked to hind limb length (relative to body size), with a ∼20% longer relative hind limb resulting in the removal of ∼30% more fire ants (Langkilde 2009). A morphological study of fence lizard museum specimens that predate the fire ant introduction revealed a latitudinal cline where lizards from the southern extreme of their range had ∼5% *smaller* relative hind limb lengths than those in the northern extreme (a difference of ∼14° in latitude), a trend likely driven by temperature and precipitation gradients (Thawley et al. 2019). However, a different pattern is observed in the present day, following this introduction. Relative limb length is greater in populations with the longest history of fire ant invasion; a pattern that could be explained by recent evolutionary change in response to fire ants (Thawley et al. 2019). A positive, albeit non-significant, correlation between mother and offspring relative limb lengths suggests that this trait is likely heritable in fence lizards (Langkilde 2009), as it is for other lizards (Tsuji et al. 1989; Kolbe and Losos 2005). Together, these observations suggest that relative limb length in fence lizards may have been and could continue to be a viable target for natural selection in response to novel interactions with fire ants.

## Results

In this study we sought to identify potential fire ant invasion-related genomic signatures of adaptation in eastern fence lizards. To do so, we analyzed whole-genome sequencing data for a total of n=420 lizards from three sampling sites (Fig 1A) – two sites were naïve to fire ants (in Arkansas, n=19, and Tennessee, n=20) and one had a long history of fire ant exposure (66 to 77 years in Alabama, n=381). We first generated high-coverage sequencing data (average of 13.01X sequence reads per site, per individual) from n=20 individuals from each site (total n=59; one sample failed QC) for population history and evolutionary genomic analyses. We also generated low-coverage (4.39X) sequencing data from an additional n=361 individuals from the fire ant-invaded site in Alabama (total n=381 when including those individuals also in the high-coverage dataset) to identify any significant genotype associations with relative hind limb length.

**Figure 1.**
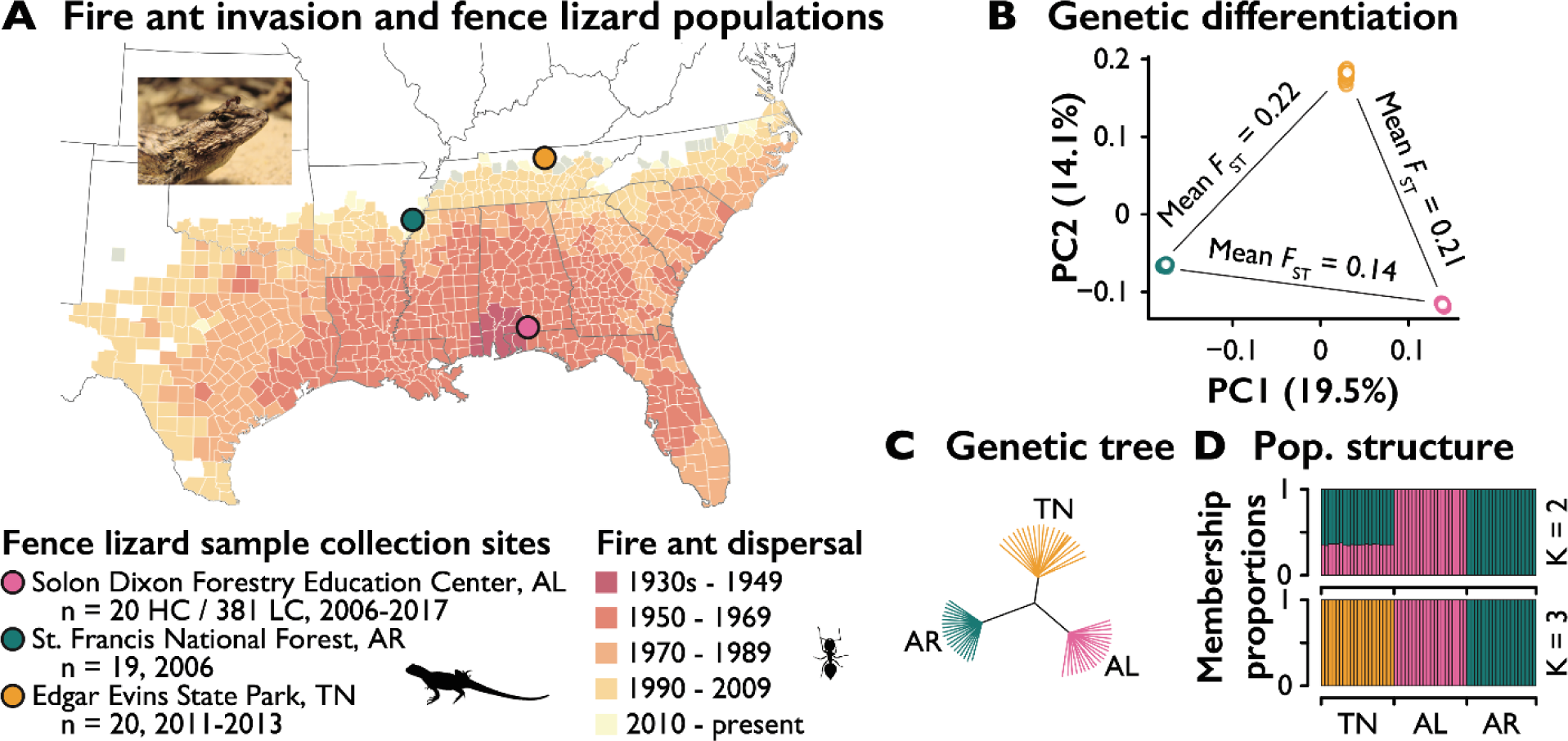
Invasive fire ant distribution and fence lizard population structure. *A*: Reported detection and quarantine of the red imported fire ant *Solenopsis invicta* in the Southeastern US since the 1930s (Code of Federal Regulations 2006), along with sampling sites, sampling periods, and number of collected eastern fence lizards, *Sceloporus undulatus*. *B.* Principal Components Analysis and mean Weir & Cockerham *F*ST values for the three pairwise population comparisons. *C*. Genetic distance-based neighbor-joining tree analysis. *D*. Admixture analysis for K = 2 and K = 3 ancestral groups. K = 3 yielded the lowest rate of cross-validation error (supp figure S2).

The lizards from the fire ant-exposed population were collected between 2006 and 2017 from the Solon Dixon Forestry Education Center, in Andalusia, Alabama (AL). Fire ants were recorded and this site was quarantined in the early 1940s (Code of Federal Regulations 2006). Estimates of eastern fence lizard generation time vary from 1.15 to 2.24 years (Rodríguez-Romero et al. 2011); thus, the approximate 70 years of coexistence with fire ants may have encompassed 31 to 60 generations of fence lizards. The other two sampled populations had no record of invasion by fire ants at the time of sampling. These lizards were collected from St. Francis National Forest, Arkansas (AR) in 2006, and from Edgar Evins State Park, Tennessee (TN) between 2011 and 2013.

We mapped sequence reads from the high-coverage dataset to the fence lizard 1.0 reference assembly (Westfall et al. 2021). After quality control and filtering (see *Methods*), we identified a total of 46,934,027 SNPs across the three populations, though only 2,249,567 of these SNPs were variable in all three populations. The AL population was the most genetically diverse (AL genome-wide mean pairwise nucleotide distance π = 0.125; TN π = 0.093; AR π = 0.076) and also had the largest number of private variants by far (AL = 24,705,862; TN = 7,360,015; AR =5,985,756). The number of SNPs variable in two of the three populations were 5,975,685 for TN and AL; 3,452,917 for AR and AL; and 4,378,018 for TN and AR. In the AL population, the presence of such a large proportion of rare genetic variants – including 10,507,067 singleton or doubleton SNPs – is consistent with a recent, large-scale demographic expansion (Slatkin 1993; Amos and Harwood 1998), even in the face of fire ant invasion.

We used several descriptive approaches to examine genetic relationships among the three populations. First, we examined patterns of population differentiation using the *F*_ST_ statistic, restricting each pairwise comparison to SNPs that were either variable in both populations or for which the minor allele in one population was fixed in the second. Average *F*_ST_ values were AL-TN = 0.210, AL-AR = 0.141, and TN-AR = 0.223 (Fig 1b). The three populations also clustered independently based on results from a principal components analysis (Fig 1b) when retaining only the 11,437,455 SNPs variable within or between at least two populations (results were qualitatively equivalent when using all SNPs; Supp fig S1). We also used this set of SNPs to construct a genetic distance matrix-based neighbor-joining tree, in which individuals from each population were distinctly separated (Fig 1c). Lastly, we used ADMIXTURE (Alexander et al. 2009) to compute model-based estimates of individual ancestry. When specifying k=3 populations (which has the lowest cross-validation error; Supp Figure S2), cluster membership proportions for members of all three populations were distinct (Figure 1d). With k=2, AL and AR membership proportions were distinct, with TN individuals exhibiting a mix of the two. Taking all these results together, despite the Mississippi River being a putative biogeographic barrier for AR fence lizards, we did not observe a clear, strong genetic separation between AR vs. AL+TN, precluding the definitive assignment of any population as a true outgroup for downstream evolutionary analyses.

### Signatures of selection exclusive to Alabama lizards overlap morphology- and immune system-related genes

To identify candidate regions for recent positive selection in each lizard population, we used three different population genetic approaches: Tajima’s D (Tajima 1989); saltilassi (DeGiorgio and Szpiech 2022), and LSBL (locus-specific branch length; (Shriver et al. 2004). If selective pressures imposed by fire ants resulted in recent genetic adaptation in the fire ant-exposed AL population, then we would predict the commensurate signatures of selection reflecting such adaptations to be absent in the two northern populations naïve to fire ants.

Tajima’s D compares the mean number of pairwise genetic differences to the sample size-corrected number of variable sites in a population across a given genomic region. Low Tajima’s D values (an excess of rare alleles) may reflect recent population expansion or positive selection (Slatkin 1993; Amos and Harwood 1998). With genome-wide deviations from D=0 most likely reflecting past demographic history, we considered genomic regions containing Tajima’s D values in the lowest 0.5% of all 100 Kb regions for a given population as positive selection candidates. For AL we identified n=143 such regions (D < −1.81; Figure 2a). For the two uninvaded populations, AR and TN, we identified n=98 (D < −2.11) and n=97 (D < −1.9) Tajima’s D candidate selection regions, respectively (Supp fig S3-S4).

**Figure 2.**
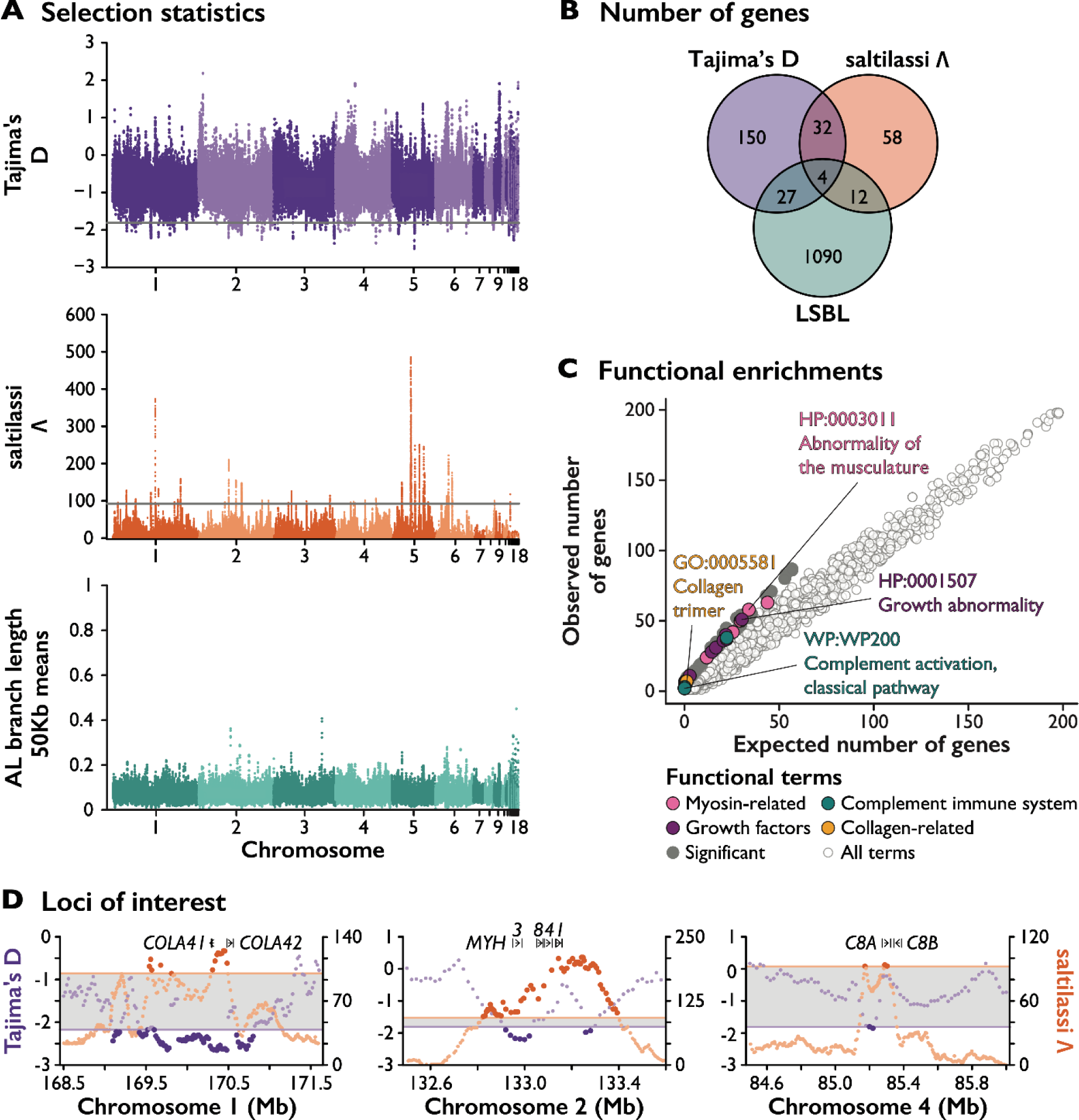
Signatures of genetic adaptation in fence lizards from the fire ant-invaded population in Alabama. *A.* Top: genome-wide Tajima’s D in 100Kb windows with a 20Kb step. Candidate selection regions fall below the 0.05^th^ percentile cutoff (D < −1.81). Middle: genome-wide saltilassi statistic for sweeping haplotypes (DeGiorgio and Szpiech 2022). Candidate selection regions fall above the 99.5^th^ percentile cutoff (Λ > 92.24). Bottom: locus-specific branch length (Shriver et al. 2004) means for the AL population in 50Kb windows and a 10Kb step. *B*. Number of genes overlapping or nearby (± 25Kb) candidate selection regions for each of the three selection statistics. *C*. Functional enrichment analysis for genes overlapping candidate regions under selection for Tajima’s D. Plausible adaptations to selective pressure from fire ants include variants in genes for myosin, collagen, growth factors, and complement immune response. *D.* Selected multi-signal candidate selection regions, with scores for the Tajima’s D and saltilassi statistics. Horizontal lines represent the same significance thresholds in *A*.

The program saltilassi computes likelihood ratio tests on the haplotype frequency spectrum of a given population to identify haplotypes with high frequencies compared to the genome-wide expectation as candidate targets of past positive selection. We similarly considered regions containing one or more variants in the top 0.5% distribution of saltilassi’s Λ statistic (see *Methods*) as selection candidate regions (for AL: Λ > 92.24 and n=1582 regions; Figure 2a; For AR: Λ > 407.56 and n = 566 regions; for TN: Λ > 252.27 and n = 764 regions; Supp Figure S5-S6).

Finally, we used the LSBL statistic (Shriver et al. 2004) to identify variants with frequencies highly differentiated in one population relative to each of the two others, based on all pairwise *F*_ST_ values (LSBL_A_ = (AB *F*_ST_ + AC *F*_ST_ – BC *F*_ST_) / 2). Here, we considered regions as candidates for positive selection if at least 3 SNPs in the top 0.1% of the LSBL distribution were observed within 50Kb of one another and in high linkage disequilibrium (r^2^ ≥ 0.9). In total, we identified n=2,210 candidate regions (LSBL > 0.76) for AL (Fig 2x), n=2,960 regions (LSBL > 0.74) for TN, and n=1,686 regions (LSBL > 0.78) for AR (Supplementary Figures S7-S8; Supplementary Table S9-S11).

We initially focused our analysis on genomic regions that were flagged as candidates for a history of past selection by at least two of the above approaches for a given population, resulting in a dataset of 42, 35, and 24 multi-signal selection candidate regions for AL, TN, and AR, respectively (Supp Table S12). We used g:Profiler (Raudvere et al. 2019) to perform functional profiling enrichment analyses to identify known biological and molecular functions and pathways (see *Methods*) significantly overrepresented among the set of genes overlapping or nearby (± 25Kb) the candidate selection regions for each population.

For the fire ant-invaded population, AL, genes within the multi-signal selection candidate regions were significantly enriched for multiple anatomical structure and the immune system functional categories (supp table S13), including Myofibril (GO:0030016; 7 observed vs. 1.21 expected genes; Fisher’s one-tailed test, FDR adjusted = 0.011) and Complement and coagulation cascades (KEGG:04610; 3 observed vs. 0.18 expected genes; FDR = 0.042). One of the multi-signal candidate regions that contains myofibril-related genes overlaps a myosin gene cluster on chromosome 2 (Figure 2d). The different myosin proteins are responsible for the distinct contractile properties across muscle cells (Weiss and Leinwand 1996; Foth et al. 2006). The complement immune system genes of interest include *C8A* and *C8B* (Figure 2d), which are directly involved in the membrane attack complex (MAC). Nucleated cells targeted by the MAC can undergo autoimmune and inflammatory processes through the secretion of proinflammatory proteins such as IL-β and IL-18 (Morgan 2016; Xie et al. 2020), making genetic variants within or nearby these genes plausible targets for natural selection by fire ant venom exposure. Lastly, another multi-selection signature (Tajima’s D and lassi) locus of interest is a 411Kb region in chromosome 1 that overlaps two genes – *COL4A1* and *COL4A2* – involved in the production of collagen IV (Figure 2). Collagen IV is the main component of the cellular basal membrane and is responsible for skeletal muscle stability (Koskinen et al. 2001; Csapo et al. 2020). In the green anole *Anolis carolinensis*, skeletal muscle is the tissue with second highest expression of *COL4A1* and the third highest expression of *COL4A2* (Bastian et al. 2021).

Among regions identified in the Tajima’s D analysis (but not necessarily as outliers with the other two statistics), growth and growth factor-related pathways were significantly overrepresented in the invaded AL lizard population (supp table S14). Significant enrichments included regulation of insulin-like growth factor transport and uptake by insulin-like growth factor binding proteins (REAC:R-MMU-381426; 7 observed vs. 1.12 expected genes; FDR < 0.01) and osteoblast signaling (WP:WP238; 2 observed vs. 0.1 expected genes; FDR = 0.042). Specific genes involved in these enrichments include the macrophage colony-stimulating factor gene *CSF1* (chromosome 4), Insulin-like growth factor 1 (*IGF1*; chromosome 5), *IGF2* (chromosome 1), and the Insulin-like Growth Factor binding protein 1 (*IGFBP1*; chromosome 6), which all serve as primary drivers of embryonic growth (Holt 2002), and fibroblast growth factor 23 (*FGF23*; chromosome 5), which has significant expression in bone tissue and is involved in osteoblast differentiation (Wang et al. 2008).

For the TN and AR populations, naïve to fire ants, genes overlapping candidate regions for positive selection in at least two statistics showed no functional enrichments analogous to those in the fire ant-invaded AL population. In TN (supp table S16), the most significant enrichment was for ovarian infertility (WP273; FDR = 0.021), while there were no significant enrichments for AR. However, when considering only the saltilassi results, there was a functional enrichment for negative regulation of developmental growth in AR (GO:0048640, 6 observed vs. 0.28 expected genes; FDR < 0.001) and analogous terms (Supp table S20).

### Genomic associations with lizard limb length in the fire ant-invaded population in Alabama

To uncover genomic associations with relative limb length variation, we brought low coverage (average 4.39X sequence coverage per individual) whole genome sequence data from an additional n=361 lizards from the same population in Alabama, into our study. We first used the high-coverage data for the 20 AL individuals (described above) to improve genotyping rates for the 361 low-coverage genomes via genomic imputation (Browning et al. 2018). Following QC and filtering, our dataset consisted of 4,245,544 SNP genotypes for n=381 total AL lizards, each with limb and body length phenotypic data available. We tested for genetic associations with relative limb length separately for each SNP using linear models, with individual sex and the first four components of a principal components analysis (to account for population structure) as covariates (see Methods). Using this procedure, we identified a total of n=24 genomic loci with significant (*P* < 1e-6) genotype-phenotype associations (Figure 3a; supp table S21; Q-Q plot available on supplemental figure S22).

**Figure 3.**
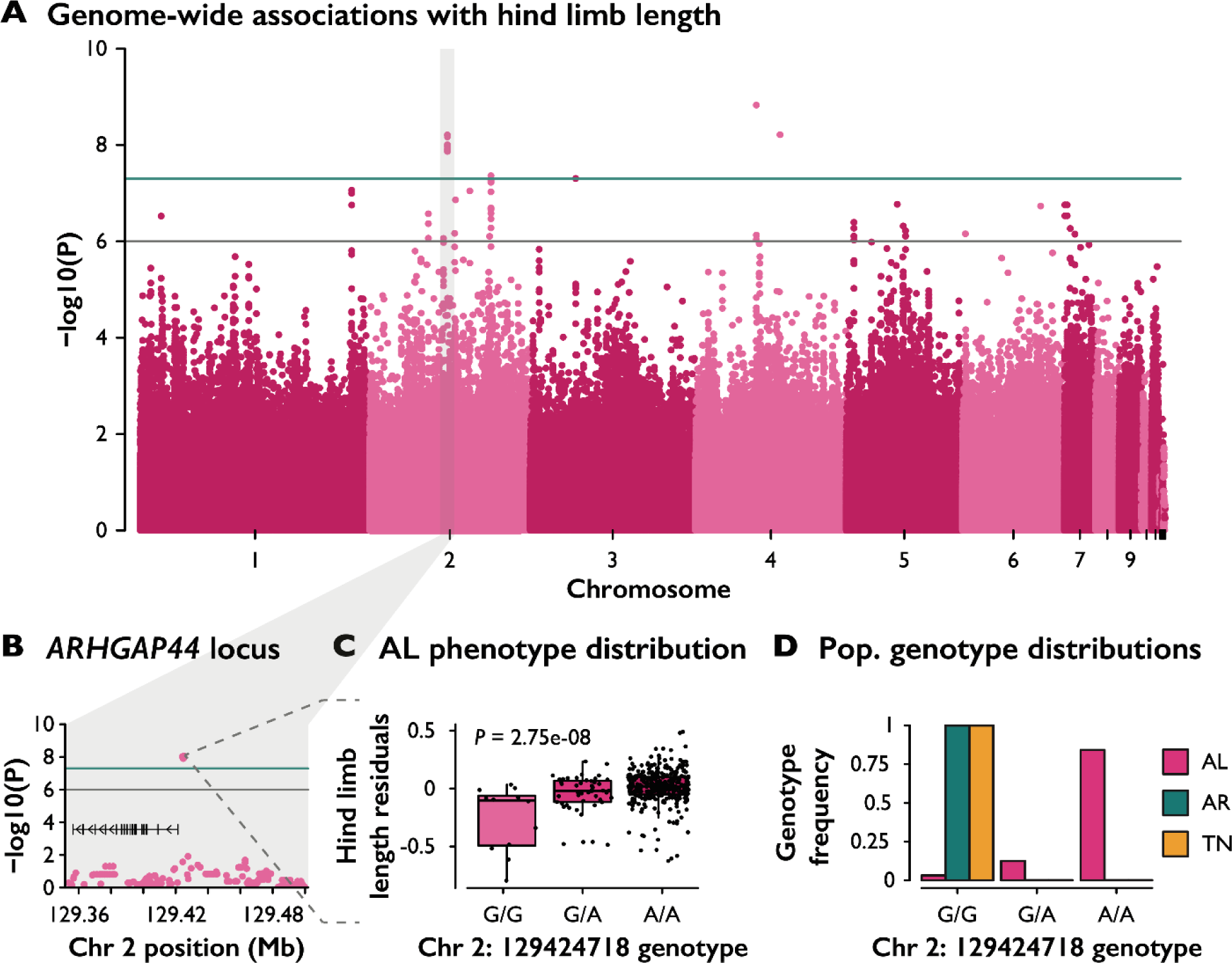
Genome-wide genotype associations with fence lizard hind limb length. *A*. Genome-wide associations with relative hind limb length (residuals of hind limb length on snout-to-vent length with sex and PCs 1-4 from a Principal Components Analysis of the genome-wide SNP genotype data as covariates). The green line indicates a significance cutoff of P < 5e-8 (four genomic loci) and the gray line a significance cutoff of P < 1e-6 (24 total loci). *B.* GWAS results for SNPs at the *ARHGAP44* gene locus, with vertical bars representing exons, and arrows representing transcriptional direction. *C*. Representative SNP immediately upstream of *ARHGAP44* and its association with relative limb length. D. For this same SNP, the negative-effect allele is fixed in the fire ant-free TN and AR populations, while the limb length positive-effect allele is abundant in the fire ant-invaded AL population.

A gene ontology enrichment analysis on the set of genes overlapping or nearby (±25Kb) these regions (supp table S23) revealed an overrepresentation of signaling by epidermal growth factor receptor pathway genes (REAC:R-MMU-177929; 2 observed vs. 0.03 expected genes; FDR = 0.049). The two genes, both members of the protein tyrosine phosphatase family, are *PTPN11* on chromosome 7 and *PTPN12* on chromosome 5. In humans, mutations in *PTPN11* have a strong association with the Noonan syndrome, characterized by short stature and skeletal malformations (Tartaglia et al. 2002).

We did not observe more overlaps than expected by chance between the GWAS regions and our AL lizard candidate positive selection regions (permutation analysis; for Tajima’s D: P = 0.31; Supp Figure S24; for saltilassi, no overlaps). However, two loci were both strongly associated with relative limb length and contained or were in proximity to alleles that were highly differentiated between AL and the fire ant-naïve AR and TN populations. The first locus overlaps *PTPN11* (chromosome 7), discussed above as part of the enrichment for signaling by epithelial growth factor receptor (Supp Fig 25). The second locus overlaps *ARHGAP44* (chromosome 2), which belongs to the gene family of Rho GTPase-activating proteins. These proteins interact with insulin-like growth factors and the CREB transcription factor to modulate body size during embryonic development (Moon and Zheng 2003). Here, three non-coding SNPs, located between 2747 and 3711 bp upstream of *ARHGAP44* exon 1 (Figure 3b), are strongly associated with relative limb length (β = 0.11, *P* = 2.75e-8; Figure 3c). For these SNPs, the alleles with positive effects on relative limb length are abundant (>84% frequency) in the AL population yet absent in TN and AR (Figure 3d). The orthologous region in humans overlaps an annotated alternative splicing isoform of *ARHGAP44* (AS-1).

## Discussion

The adaptive potential of natural populations when faced with the sudden and pervasive impacts of anthropogenic activity is uncertain. Understanding these fundamental drivers of ecosystem change is crucial—especially in an era of unprecedented human activity and climate change. Our study focused on endemic fence lizards, who have co-existed with introduced fire ants for more than 70 years in parts of the Southeastern United States. We sampled lizards from a long-invaded site in Alabama and identified multiple genomic signatures of positive selection that were generally absent from two lizard populations naïve to fire ants, in Arkansas and Tennessee.

### Putative adaptations to fire ants

The set of genes contained within or nearby the candidate signatures of positive selection that we identified in lizards from the fire ant-invaded AL population were significantly enriched for morphological development functions. Two additional loci – which respectively overlap and are immediately upstream of genes involved in body growth processes – contain alleles that are simultaneously strongly associated with longer legs in AL lizards and occur in regions of high genetic differentiation relative to the fire ant-unexposed TN and AR populations. These variants may constitute some of the genetic adaptations underlying fence lizard limb length variation, with longer hind limbs associated with increased survival against swarming ants (Langkilde 2009).

We also identified multiple signatures of positive selection for variants within or nearby two genes associated with membrane attack complex inflammatory processes (Morgan 2016; Xie et al. 2020). Inflammatory processes are a key component of the immunological response to cellular damage from venom toxins (Stafford 1996; dos Santos Pinto et al. 2012; Zamith-Miranda et al. 2018; Liu et al. 2023). Remarkably, *S. undulatus* field and laboratory experiments have previously detected associations between fire ant venom and complement immune activity (Tylan et al. 2020; Tylan et al. 2023), which includes the membrane attack complex. Specifically, field-caught lizards from AL had significantly lower levels of complement activity relative to northern, unexposed lizards (Tylan et al. 2020). Separately, lizards from a TN population naïve to fire ants showed higher levels of complement activity immediately after consuming fire ants, as well as three weeks post fire ant stinging (Tylan et al. 2023). Therefore, it seems likely that the complement immune system is involved in both routes of exposure to fire ant venom, albeit with temporally distinct signals for each.

Given this context, the findings from our study lead us to suggest that population-level differences in complement immune activity may in part be a product of recent evolutionary processes in response to the introduced fire ant. Specifically, we hypothesize that variants within the *C8A* and *C8B* membrane attack genes candidate selection region that we detected in the fire ant-exposed AL population but not the fire ant-naïve TN and AR populations may underlie an adaptive suppression of the innate complement system. If constant consumption of fire ants triggers chronic, costly inflammatory responses that are of limited immunological benefit, then it follows that such inflammatory suppression will be sparing of resources otherwise diverted from other physiological processes (Straub and Schradin 2016). Such a tradeoff between immune response and energetic resources has been demonstrated in poultry (Van Der Most et al. 2011), house sparrows (Martin et al. 2003), and the tobacco hornworm (Adamo et al. 2017).

### Community-scale consequences

Across ecosystems, evidence for rapid adaptation to human-induced environmental changes continues to build (Strauss et al. 2006; Shaw and Etterson 2012; Sullivan et al. 2017; McCulloch and Waters 2023). A few illustrative examples include the water flea *Daphnia* showing signals of adaptation for resistance to salinization derived from human activity (Wersebe and Weider 2023), and independent populations of crested anoles (*Anolis cristatellus*) carrying signatures of convergent genetic adaptation to urbanization (Winchell et al. 2023). Meanwhile, human-commensal house sparrows (*Passer domesticus*) have signatures of positive selection overlapping an amylase gene, which encodes an enzyme involved in starch digestion (Ravinet et al. 2018), and poaching of African elephants for ivory appears to have resulted in rapid adaptation for a tuskless phenotype (Campbell-Staton et al. 2021).

Although the above examples and our results in fence lizards following the introduction of fire ants contribute to the growing evidence of successful native species adaptations in response to anthropogenic change the species-specific and community-wide ramifications of rapid adaptation in a keystone species remain unknown (Thompson 1999; Mooney and Cleland 2001; Strauss et al. 2006). For fence lizards, while fire ant predation is at least partly counteracted by longer hind limbs combined with a twitch/flee response for fire ant removal and escape, such behavior is likely accompanied by a break in crypsis. Field surveys in an Alabama fence lizard population have demonstrated that close to 50% of male lizards show signs of injuries – a two-fold increase from fire ant-naïve populations – speculatively due to increased detection by birds of prey (Thawley and Langkilde 2017). This conundrum likely represents a delicate balance between context-dependent antipredator responses (Martín et al. 2009) faced by fence lizards since fire ant introduction, exemplifying the broader community ramifications of adaptation to human activity.

Meanwhile, if it is true that fence lizards from fire ant invaded sites, such as in Alabama, have adapted to exploit fire ants as a novel food resource, as our results suggest, then this may have significant cascading effects on that ecosystem’s food web. The voracious predatory behavior of fire ants has been broadly demonstrated; they can effectively prey on higher-level consumers such as salamanders (Todd et al. 2008), bird nestlings (Kopachena et al. 2000), cotton rats (Long et al. 2015), and hatchling sea turtles (Allen et al. 2001). Consequently, with potential adaptations to include fire ants in their dietary niche, fence lizards may indirectly assimilate biomass otherwise unavailable. The ability to exploit this nutritional input may be one factor underlying a recent, rapid population expansion of AL fence lizards, evidenced by an abundance of rare alleles observed in our analysis. Such food web disruptions by invasive species are to be expected (Zanden et al. 1999; Miehls et al. 2009). However, less discussed is how standing genetic variation can capitalize on rapid anthropogenic change for an ecological advantage.

In conclusion, our study identifies genetic signatures of positive selection in fence lizards exposed to human-introduced fire ants. These plausible adaptations to the fire ant introduction are observed in conjunction with a recent and large-scale population expansion in fence lizards, inferred from our population genomic data. Together, our findings highlight the potential of standing genetic variation in promoting population resilience in the face of anthropogenic disturbance.

## Methods

### Animal sampling

Between 2006 and 2017, adult male and female lizards were collected from the three study populations using the lasso method. The AL population was included in the study due to its long history with introduced fire ants. The individuals from the TN and AR populations were included because these sampling locations had not yet been invaded by fire ants at the time of collection. Phenotypic data were collected from the n=381 sexually mature lizards in our AL population sample: hind limb length (HLL) and snout-to-vent length (SVL) were each measured to the nearest 0.5 mm using a ruler following a protocol described in (Langkilde 2009) The phenotypic data are available in supp table S26.

### DNA Extraction

All the fence lizard tissue toe and/or tail samples were stored in 70% ethanol at 4°C. We used up to 30 mg of each of the 421 preserved fence lizard tissue samples for E.Z.N.A.® tissue kit (D3396, Omega Bio-Tek, Inc., Norcross, GA, USA) DNA extractions. DNA extractions were performed following the manufacturer’s instructions with the following exceptions: each tissue sample was ground with a polypropylene pestle in a 1.5-mL microcentrifuge tube, total digestion time was increased to 14-15 hours in a 600 rpm shaking thermomixer, 1 μL of Pellet Paint® NF Co-Precipitant was added to each sample to increase DNA adherence in the HiBind® DNA Mini Column, and the total elution volume was halved (two 50-μL portions). Each sample’s DNA extraction concentration was obtained with a Qubit® 3.0 Fluorometer dsDNA High Sensitivity Assay Kit, and then stored at −20°C until library preparation.

### Library Preparation and Sequencing

Portions of each DNA extract were sheared to a target length of 500 bp with a Covaris M220 Focused-ultra sonicator (Peak Incident Power: 50, Duty Factor: 20%, Cycles per Burst: 200, Temperature: 20°C). Libraries for each sample were prepared from ≥200 ng of sheared DNA with TruSeq® Nano DNA High Throughput Library Prep Kit (20015965, Illumina Inc., San Diego, CA, USA) and IDT for Illumina – TruSeq® DNA UD Indexes v1 (Illumina Inc., San Diego, CA, USA). The libraries were pooled and sequenced with a paired-end 150 bp strategy on two Illumina NovaSeq 6000 S4 flowcells for 1.3 T of paired-end raw read data each. One pool (HC-60) had 20 randomly selected AL individuals as well as the 20 lizards from the uninvaded sites TN and AR. An average of 165.24 million reads were generated for each sample in pool HC-60. The second pool (LC-381) contained the full set of 381 AL individuals, with an average 27.88 million reads sequenced.

### Read Mapping and Quality Filtering

We used a chromosome-level reference genome assembly, “PBJelly,” that was recently developed from two male *S. undulatus* individuals collected at Solon Dixon Forestry Education Center, AL (English et al. 2012; Westfall et al. 2021)for read mapping. The annotated reference assembly was indexed with bwa v0.7.16 index and SAMtools v1.5 faidx (Li et al. 2009; Li and Durbin 2009), and a sequence dictionary was created with picard CreateSequenceDictionary (Picard Toolkit 2019) for use in read mapping, SNP identification, and downstream analyses.

The LC-381 group reads were sequenced without lanes in their NovaSeq S4 flowcell, but the HC-60 group reads were sequenced across four lanes and needed to be combined into one forward and reverse read prior to trimming and mapping to the reference genome. The raw reads were trimmed with Trimmomatic v0.36 to remove the Illumina TruSeq3-PE-2 adapters and other reads <36 bases long, as well as leading and trailing low quality or N bases (Bolger et al. 2014). The trimmed reads were aligned to the reference genome with bwa v0.7.16 mem (default settings), an alignment tool specialized for large genome sizes that seeds alignments with maximal exact matches and extends seeds with Smith-Waterman’s affine-gap penalty for insertions or deletions (Li 2013).

SAMtools v1.5 flagstat was used to calculate estimated genome-wide coverage for the mapped reads, and “-view” was used to convert the mapped .sam files to .bam files (Li et al. 2009). SAMtools BAMtools v2.4.1 was used to sort and filter out unmapped reads and mapped reads with mapQuality less than 50 (Li et al. 2009; Barnett et al. 2011). The subset of AL (Solon Dixon) samples subject to both high- and low-coverage sequencing (n=20) were merged after mapping (using samtools -merge) and were processed as part of the LC-381 pool. Duplicates were marked in all samples (AL, TN, AR) using Picard MarkDuplicates (Picard Toolkit 2019). Read groups were added to the mapped read files with Picard AddOrReplaceReadGroups (Picard Toolkit 2019), then the reads were sorted and indexed with SAMtools v1.5 (Li et al. 2009). Sequence metrics were collected using Picard CollectWgsMetrics (Picard Toolkit 2019). For the remaining analyses, we found that several of the more computationally intensive programs required working iteratively at the chromosome-rather than whole-genome level to finish processing and within the limits postulated by our computational cluster system; we indicate such cases accordingly.

### SNP Identification

We followed the Genome Analysis Toolkit (GATK, v4.2.0.0) “Best Practices” pipeline for germline short variant discovery in each of the sequencing pools (McKenna et al. 2010; DePristo et al. 2011; der Auwera et al. 2013). Even though GATK’s pipeline was designed and optimized for analyzing human genetic data, it has been successfully applied in multiple non-model systems with available high-quality reference genomes for evolutionary genomic inferences (Kryvokhyzha et al. 2019; Wright et al. 2019; Bernhardsson et al. 2020; Chen et al. 2020; Wang et al. 2020) and it has been reported to outperform other variant callers in capability and accuracy (Liu et al. 2013; Pirooznia et al. 2014).

GATK’s pipeline began with HaplotypeCaller calling germline SNPs and indels for each individual via local *de novo* assembly. In short, HaplotypeCaller defined active regions based on the presence of evidence for allele variation in each individual’s mapped reads, then built a De Bruijn-like graph to detect overlaps between sequences and reassemble the active region (Poplin et al. 2018). The possible active regions were realigned against the reference haplotype with the Smith-Waterman algorithm to identify potential variant sites, i.e. single nucleotide polymorphisms (SNPs; (Poplin et al. 2018). Likelihoods of alleles were determined using GATK’s PairHMM algorithm, and the most likely genotype per Bayes’ rule was assigned to each potentially variant site.

HaplotypeCaller generated an intermediate GVCF file that contained likelihood data for every position in each of the top 24 largest chromosomes in every individual’s mapped read data. The per-chromosome GVCFs were merged and then indexed then merged GVCF files with GATK’s IndexFeatureFile program. Following the “Best Practices” pipeline,

GenomicsDBImport was used to import the single-sample GVCFs into a per-chromosome database (GenomicsDB) before joint genotyping with GenotypeGVCFs (per-chromosome, subset into six intervals each for the largest chromosomes, scaffolds 1-6). The resulting chromosome VCFs were then combined with VCFtools v0.1.12 “vcf-concat” (Danecek et al. 2011) into one file for each sequencing pool. There were 59,006,281 possible SNP sites identified in the HC-60 group and 67,124,902 SNP sites in the LC-381 group (bcftools -stats; bcftools v.1.12; (Danecek et al. 2011)

### SNP Filtering and Quality Control

The raw SNPs were filtered with a series of thresholds recommended by GATK (Poplin et al. 2017). GATK SelectVariants kept only variants that were classified as SNPs, then VariantFiltration removed SNPs with hard-filters based on the INFO and FORMAT fields of the VCF files: quality score by depth (QD) <2.0, Phred-scaled p-value using Fisher’s exact test (FS) >60.0, and mapping quality score (MQ) < 40.0. SelectVariants was applied again to only keep SNPs that were not filtered out by VariantFiltration. After GATK filtering there remained 56,598,888 SNPs in the HC-60 group and 64,214,575 SNPs in the LC-381 group.

The SNPs that remained in each pool after GATK’s suggested parameters were additionally filtered with VCFtools v0.1.12 for analysis (Danecek et al. 2011) Both pools were filtered to keep only biallelic sites (min-alleles 2, max-alleles 2) and remove sites with insertions and deletions (Danecek et al. 2011). The HC-60 pool was also filtered for Hardy-Weinberg Equilibrium with a low enough setting to remove sites that were likely to be erroneous variant calls (hwe 0.000001), leaving 49,837,059 SNPs for analysis (Danecek et al. 2011). PLINK (v1.9) was also used to filter the HC-60 VCF with -geno 0.05 and -mind 0.1 flags to filter out variants with missing call rates prior to downstream population analyses, with 46,934,026 SNPs remaining (Chang et al. 2015). The LC-381 group was also filtered for Hardy-Weinberg Equilibrium (hwe 0.001) and to remove sites with a minor allele frequency (MAF, number of times an allele appears over all individuals at that site divided by the total number of non-missing alleles at that site) less than 0.05 to prevent inflation in downstream statistical estimates and during imputation with the remaining 458,533 SNPs (Danecek et al. 2011).

To improve genotyping rates for the LC dataset, we leveraged the 20 HC sequences from AL as a template for genomic imputation. To that end, we first used Shape-IT version 2.r837 (Delaneau et al. 2008) to phase the each of the 24 scaffolds of the 20 HC sequences and obtain haplotype files. These were used as reference for the imputation of 381 LC sequences. Imputation was performed with Beagle 5.2 (Browning et al. 2018)using a window size of 100 and overlap of 10. Prior to the genome-wide genotype-phenotype association analysis that was conducted with the n=381 AL lizard dataset, we removed genotypes with minor allele frequencies < 0.05, leaving 4,245,544 SNPs.

### Demographics analyses

We first calculated the mean genome-wide *F*_ST_ between each of the three pairs of populations using the VCFtools function --weir-fst-pop (Danecek et al. 2011). For this analysis, we filtered the SNP dataset to remove any SNPs whereby an allele that is fixed in one population is also the minor allele in the second population (for AL/TN, SNP count n = 16,075,146; for AL/AR, n = 33,225,824; for TN/AR, n = 16,383,584). We then ran three demographics analyses involving the three populations: a principal components analysis (PCA), admixture analysis, and a genetic neighbor-joining tree based on a genetic distances matrix. For these, the original set of 46,934,026 SNPs was filtered to remove any SNPs whereby an allele that is fixed in two populations is also the minor allele in the third population, leaving 11,437,455 SNPs. The PCA was calculated using PLINK’s –pca function (Chang et al. 2015). Population structure was inferred using ADMIXTURE (Alexander et al. 2009) for a number of ancestral populations K of 2 through 5. The neighbor-joining tree was built using the R package phangorn (Schliep 2011) with a sample genetic distance matrix generated with PLINK (Chang et al. 2015).

### Genome-Wide Estimates of Tajima’s D

For each population, Tajima’s D was estimated in windows across the genomes using vcf-kit v0.2.6 (Cook and Andersen 2017) Windows were 100 Kb in length with a 20 Kb step. For each population, putative windows under recent positive selection were those in the bottom 0.5^th^ percentile of the genome-wide distribution (*i.e.*, D < −1.81 for AL; D < −2.11 for AR; and D < −1.9 for TN).

### Genome-wide Haplotype Distributions via saltilassi statistic

The filtered VCF files were assessed for signatures of positive selection with lassip v1.1.1 using the “saltiLassi” method (DeGiorgio and Szpiech 2022) The VCF files for each population were further separated into a single VCF file for each scaffold. A population ID file was created containing each individual sample ID and the corresponding population IDs. Each scaffold was passed to lassip with the following parameters to calculate haplotype statistics and the haplotype frequency spectrum (HFS): --hapstats –winsize 201 –k 20 –calc-spec –winstep 100. The genome wide average of the HFS, which functions as the null spectrum, was then determined before calculating the saltilassi statistic, a likelihood ratio statistic denoted as Λ, for each chromosome. For each population, haplotypes in the top 0.5% of the genome-wide Λ (*i.e.*, Λ > 92.24 for AL; Λ > 407.56 for AR; and Λ > 252.27 for TN) distribution were designated as candidate signatures of positive selection.

### Locus-specific Branch Lengths (LSBL)

We used LSBL to identify genomic regions in the AL population that are significantly differentiated from the two northern populations I to fire ants. We first used VCFtools to calculate the per-SNP *F_ST_* of the three pairwise population comparisons, and then used those values to calculate LSBL for AL as per Shriver *et al*. (2004): LSBL = (AL-AR *F_ST_* + AL-TN *F_ST_* – TN-AR *F_ST_*). We identified SNPs in the top 0.1% of the genome-wide distribution (LSBL > 0.76) and combined these outlying SNPs into regions that grouped all outlying SNPs within 50 Kb and in high linkage disequilibrium (LD) with each other (R^2^ >= 0.9) into a single region. LD was calculated using vcftools (--geno-r2) (Danecek et al. 2011). Next, we used the R package *ivs* (Vaughan 2023) to combine linked SNPs into candidate genomic regions.

### Functional profiling

For each population, we investigated functional enrichments in the sets of genes located within or nearby (±25 Kb) candidate regions for positive selection identified by any two of the three selection statistics (Tajima’s D, saltilassi, or LSBL; supp table S12), and separately for Tajima’s D only and saltilassi only, for each population. To that end, we used the g:GOSt function of the g:Profiler platform (Raudvere et al. 2019) based on functional annotations for *Mus musculus* and across all available databases (*i.e.*, GO: molecular function, GO: biological process, GO: cellular component, KEGG, Reactome, WikiPathways, TRANSFAC, miRTarBase, Human Protein Atlas, CORUM, HP). The background set of genes were all genes in the SceUnd 1.0 assembly (Westfall et al. 2021). We corrected for multiple tests using the False Discovery Rate (FDR; Benjamini and Hochberg 1995).

### Genome-wide Association Study

Relative limb length was calculated by extracting the residuals of a linear regression for HLL on SVL. The residuals were then assigned as the response variable for all models. We used the imputed genotype dataset (see *SNP Filtering and Quality Control* above) for the GWAS. Each SNP was numerically coded for each biallelic genotype (0, 1, or 2) and included as a predictor in its respective model. Covariates were individual sex, to control for sex-specific morphological differences, along with the eigenvectors of a population principal components analysis of the unimputed LC genomic dataset, to help control for population structure (Supp figure S27). Principal components 1 through 4 were included in each model, which in combination explained a total of 23.24% variation explained. Results for each of the 4,245,544 models are available in supp table S21.

### Permutation Analysis

To test whether candidate regions for positive selection and genomic regions associated with limb length overlapped each other more often than expected by chance, we performed permutation analyses with the R package *regioneR* (Gel et al. 2016). For each pairwise permutation (e.g., pairwise comparisons between GWAS and Tajima’s D, saltilassi, LSBL, and the set of regions highlighted by at least two out of the three statistics), the set of regions for one statistic was held static while regions of same length as those from the second statistic were randomly placed across the genome in each of 10,000 permutations. To constrain the available genomic space wherein regions were to be permuted, the number and length of chromosomes was specified using the argument *genome* in the *permTest* function. These random permutations generated a neutral probability distribution, and we assessed whether our observed number of overlaps significantly deviated from this distribution (α < 0.05).

## Supporting information

Appendix III

## Data availability

Raw sequence data have been deposited in NCBI SRA BioProject: PRJNA656311. Quality-controlled and filtered VCF’s and the full outputs for the GWAS and evolutionary genetics statistical analysis are available on Dryad https://doi.org/10.5061/dryad.tht76hf50. Scripts for all statistical analyses are available on https://github.com/braulioassis/sce-sol.

## Acknowledgements

Animal collections were approved by the Pennsylvania State University Animal Care and Use Committee, and by the respective States where collection occurred. We thank the Lansdale family for allowing us to collect lizard individuals on their property. We also thank T. Schwartz and the *Sceloporus* Genome Collaboration team, T. Adams for assistance in the curation of lizard specimens, and C. Roberts and K. Hickmann for assistance in laboratory work. We thank L. Perez and N. Grube for bioinformatic support, and S. Giery and J. Schluter for comments on the manuscript. We also acknowledge the NYU Langone Genome Technology Center (RRID: SCR_017929). This shared resource is partially supported by the Cancer Center Support Grant P30CA016087 at the Laura and Isaac Perlmutter Cancer Center. Computations for this research were performed on the Pennsylvania State University’s Institute for Computational and Data Sciences’ Roar supercomputer. Components of this work were supported by the National Science Foundation Graduate Research Fellowship Program (DGE-1255832, to A.P.S.), the National Science Foundation (BCS-1554834, to G.H.P), and the Penn State University College of the Liberal Arts.

