## Appendix I for "Genomic signatures of adaptation in native lizards exposed to human-introduced fire ants"

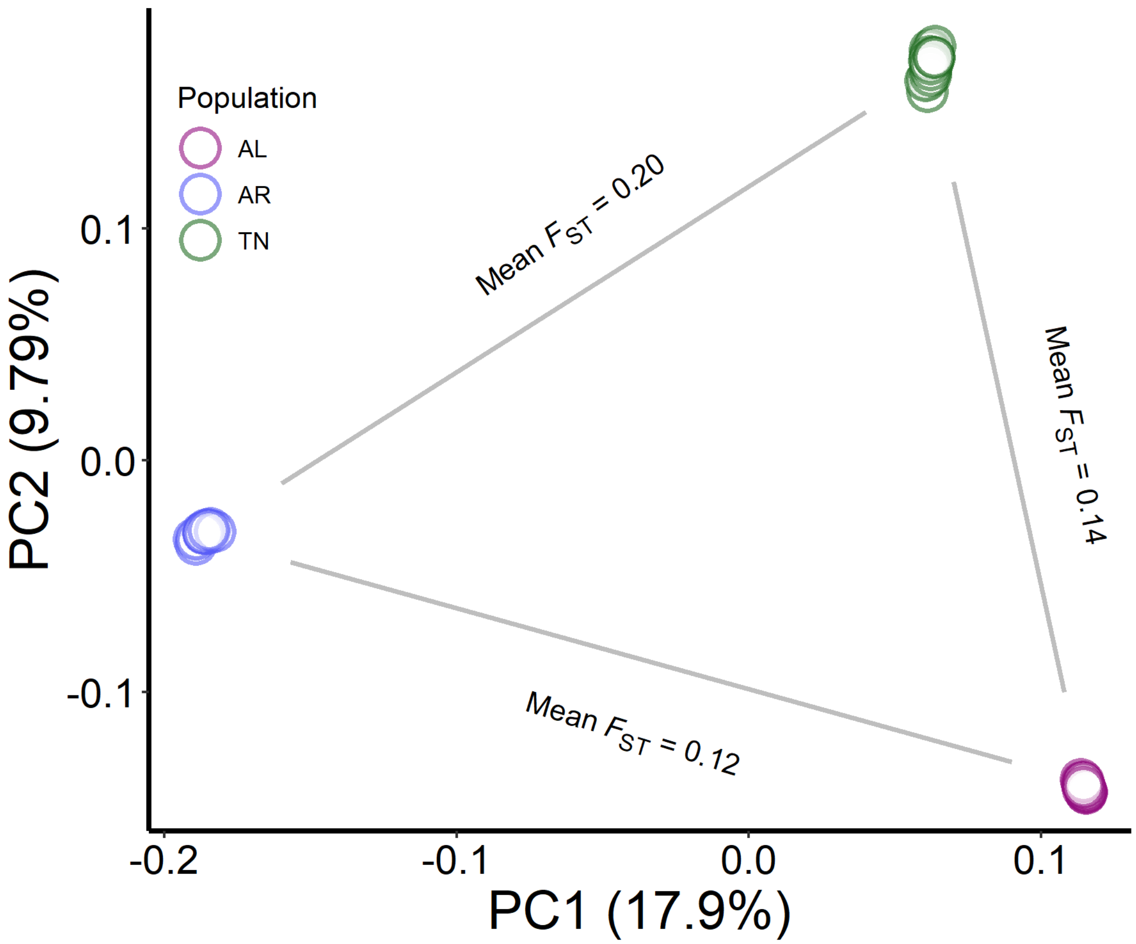


Fig S1: Principal Components Analysis and mean Weir & Cockerham $F_{ST}$ for the three pairwise population comparisons based on the full set of 46,934,027 filtered SNPs (see *Methods*).

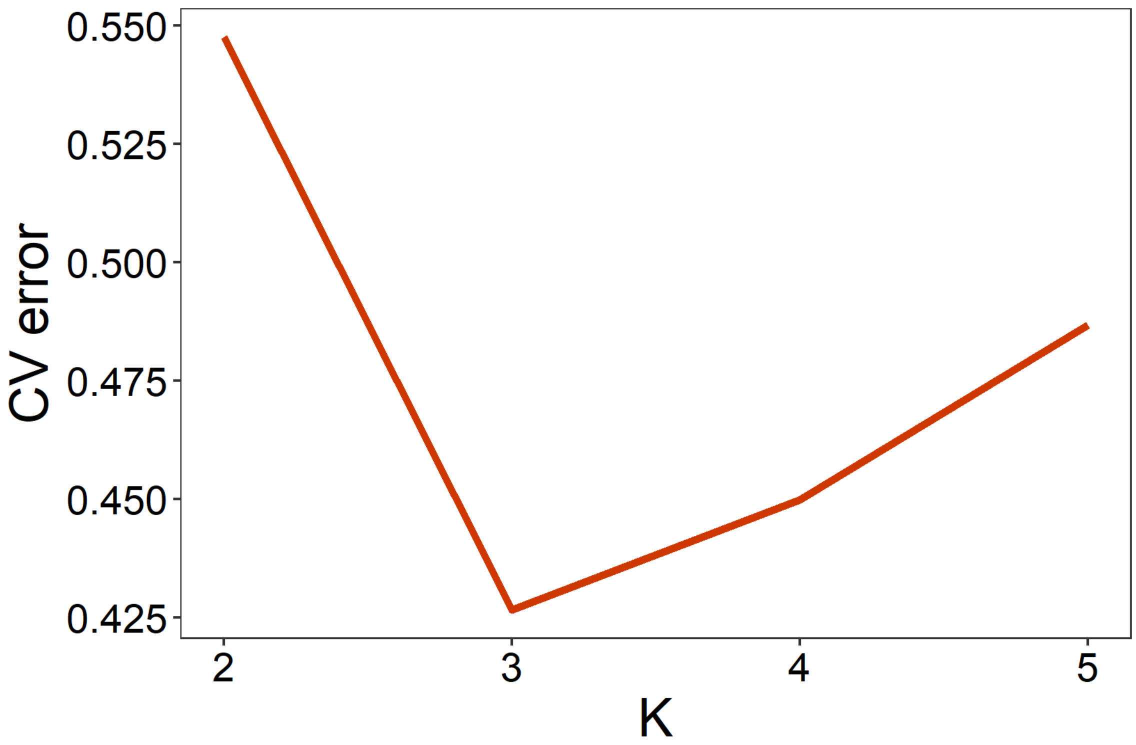


Fig S2: Cross-validation errors for admixture analysis with for K=1 through K=5 specified ancestral populations.

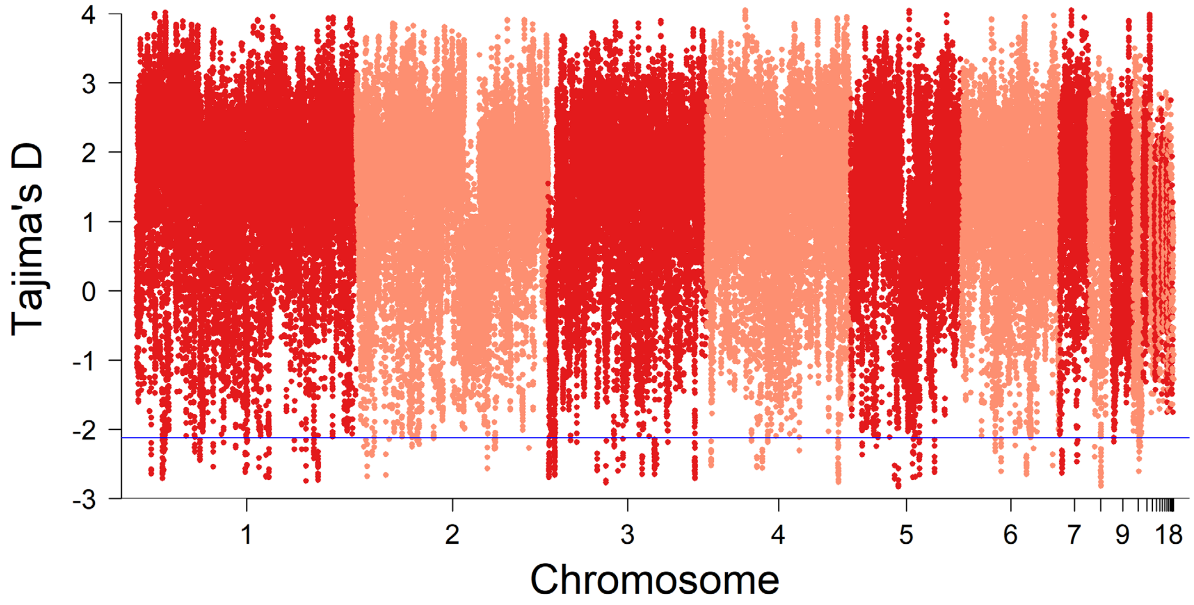

Fig S3: Genome-wide Tajima's D for the Arkansas population in 100 Kb windows and a 20 Kb step. Windows significantly under positive selection fall below the 0.5$^{th}$ percentile of the genome-wide distribution (D < -2.11, blue line).

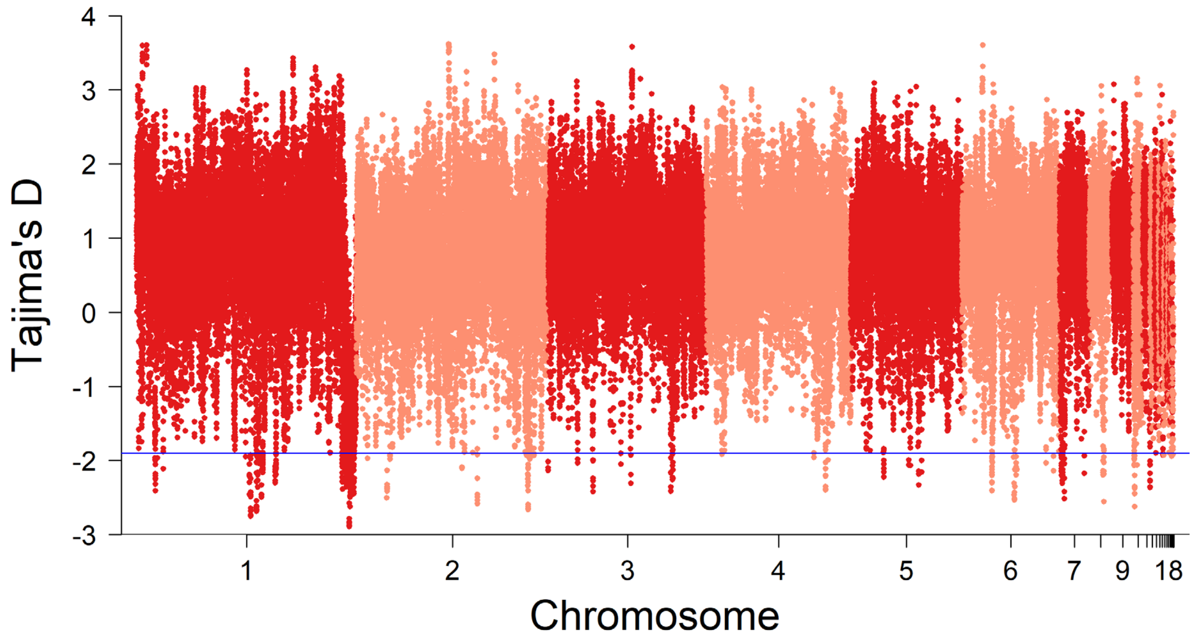


Fig S4: Genome-wide Tajima's D for the Tennessee population in 100 Kb windows and a 20 Kb step. Windows significantly under positive selection fall below the 0.5$^{th}$ percentile of the genome-wide distribution (D < -1.9, blue line).

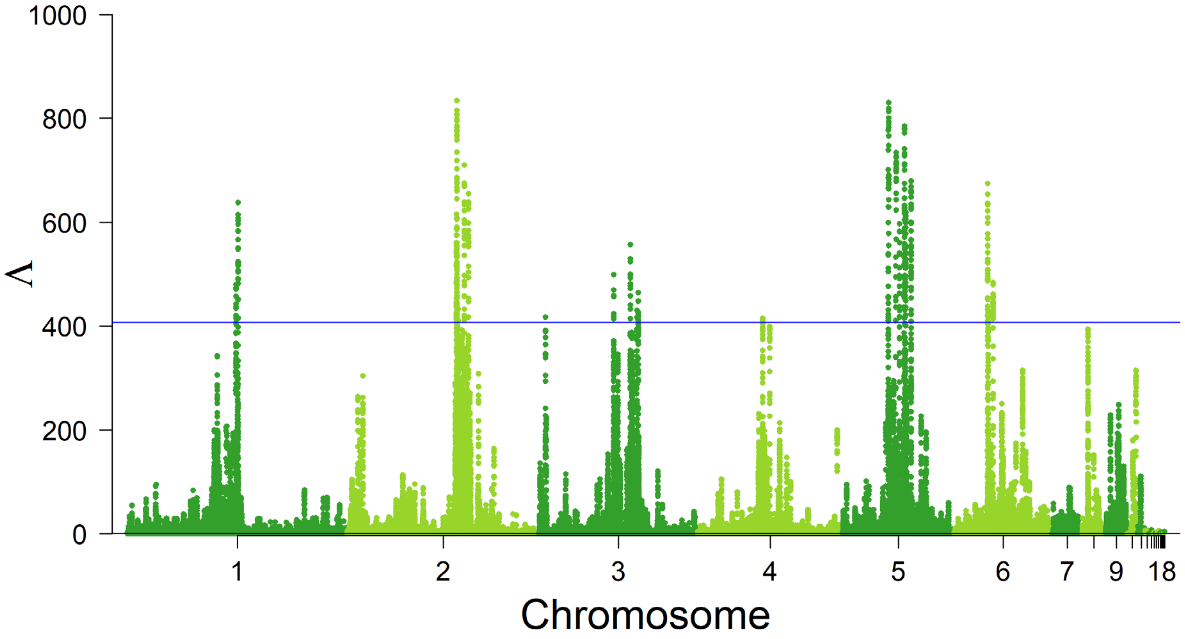


Fig S5: Genome-wide saltilassi's Λ statistic for the Arkansas population. Haplotypes significantly under positive selection fall above the 99.5$^{th}$ percentile of the genome-wide distribution (Λ > 407.56, blue line).

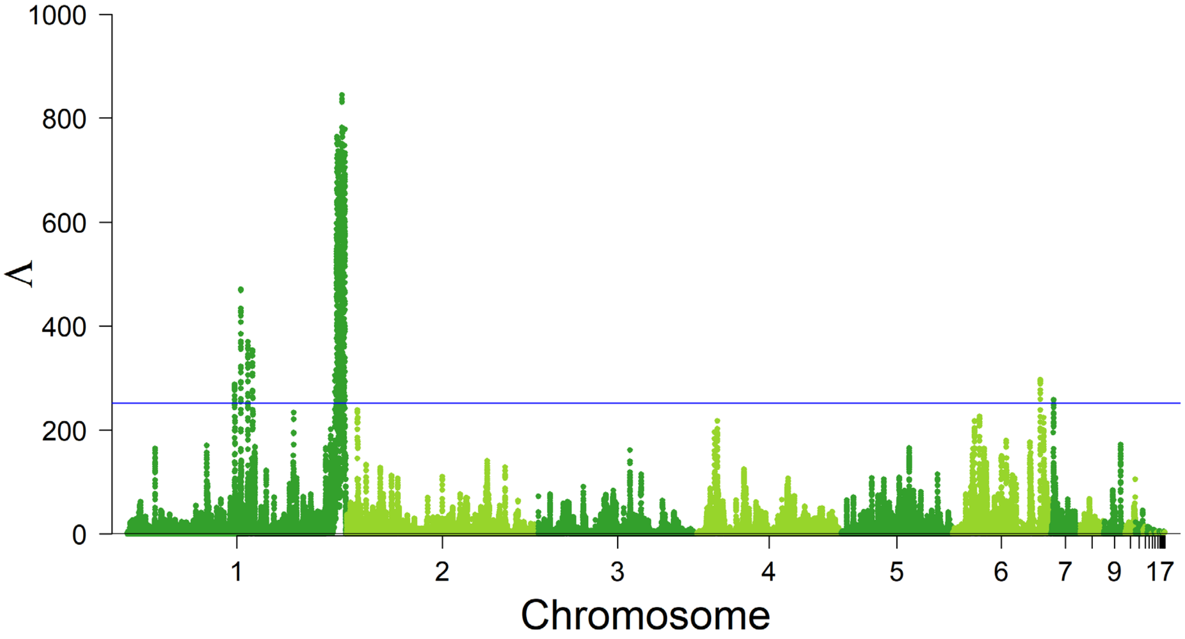


Fig S6: Genome-wide saltilassi's Λ statistic for the Tennessee population. Haplotypes significantly under positive selection fall above the 99.5th percentile of the genome-wide distribution (Λ > 252.27, blue line).

0

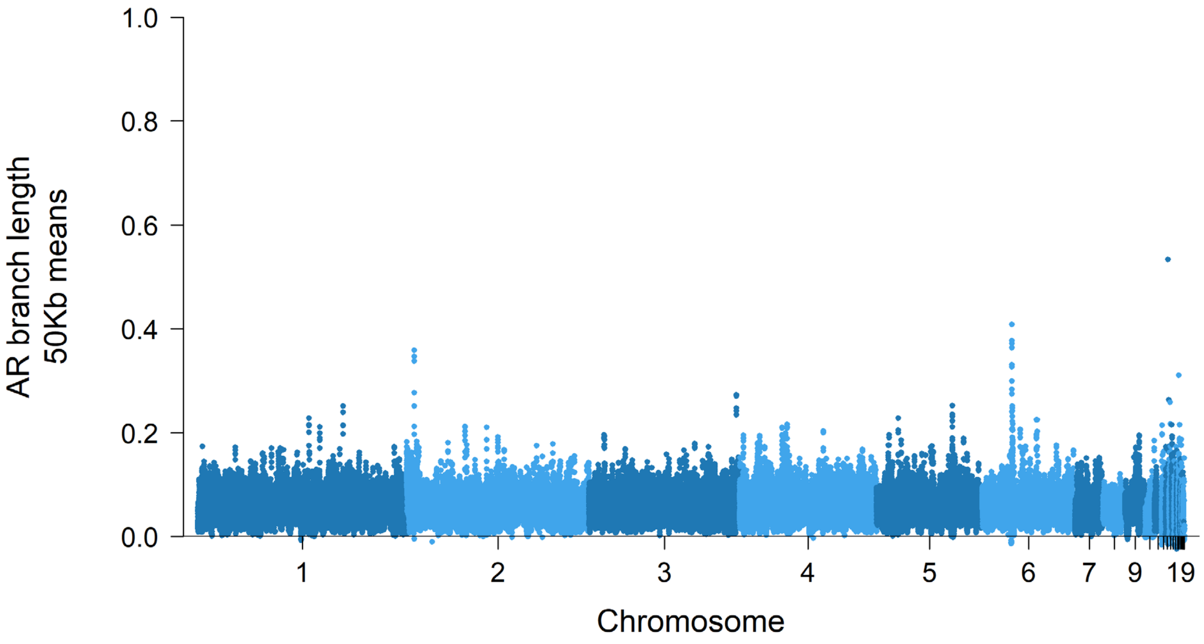

Figure S7: Locus-specific branch length for the Arkansas population (AR-AL $F_{ST}$ + AR-TN $F_{ST}$ – TN-AL $F_{ST}$) in 50Kb means and a 10Kb step.

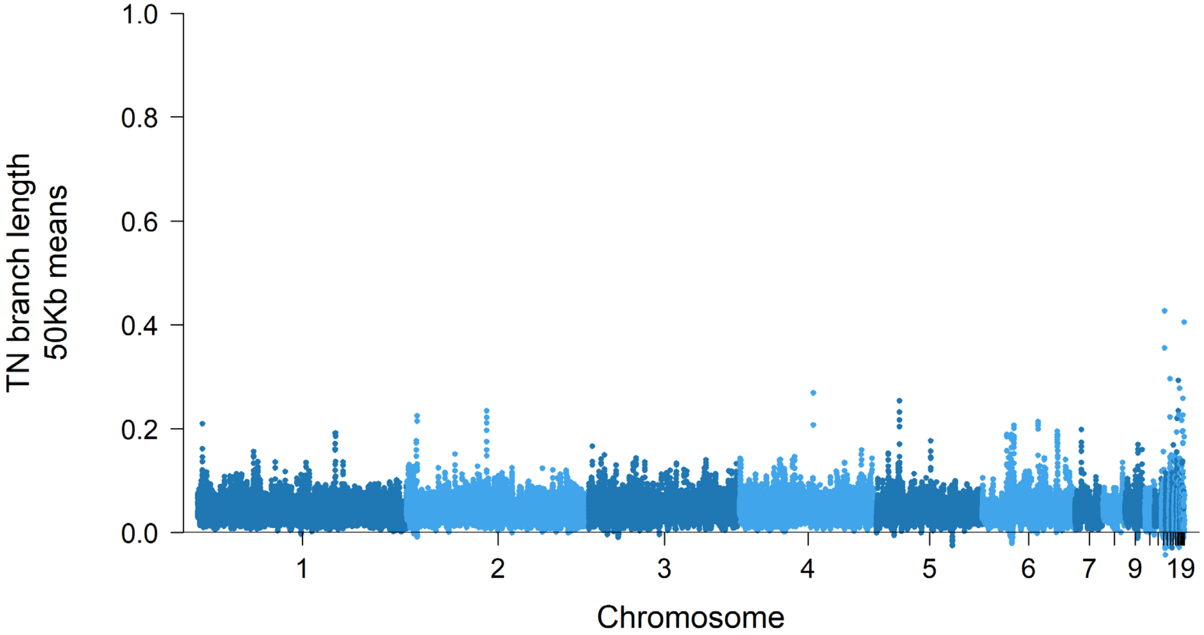


Figure S8: Locus-specific branch length for the Tennessee population (TN-AL $F_{ST}$ + TN-AR $F_{ST}$ – AR-AL $F_{ST}$) in 50Kb means and a 10Kb step.
